# Bumble bees that follow a stricter routine innovate less: Foraging behaviors, environmental complexity, and how they relate to novel problem solving

**DOI:** 10.64898/2026.03.06.710156

**Authors:** Shannon R. McWaters, James J. Kearsley, David W. Kikuchi, Timothy J. Polnaszek, Anna Dornhaus

## Abstract

The ability of animals to innovate - solve novel problems - can shape their ecology and evolution. Here we investigate how individual traits and environmental complexity relate to successful solving of a novel problem. We presented foraging bumble bees (*Bombus impatiens*) with artificial flowers of not-previously-encountered shapes and recorded the bees’ latency to access nectar. We measured individual foraging traits across multiple trips with simple flowers that did not require innovation, and bees were foraging either in a simple or complex environment (cluttered flight arena). Bees in complex environments took longer to find and were less likely to land on novel flowers, indicating that environmental complexity may take up cognitive resources and make search more difficult. However, we did not find an effect of environmental treatment on the ability or time to access reward in novel flowers once bees had landed on them. In contrast, behavioral traits significantly predicted how quickly bees ‘solved’ novel flowers. In particular, overall foraging tempo as well as routine formation, i.e. how much bees followed a fixed route on known flowers, predicted innovation - faster bees innovated faster, and bees with more repetitive foraging sequences were slower to solve the novel tasks.

Overall, while the degree of evolutionary ‘novelty’ in tasks or solutions is always hard to evaluate, our findings demonstrate that environment and individual traits may affect innovation in different ways. Individuals in simple environments may be more likely to *detect*, and individuals that are generally faster and have a lower tendency to develop fixed routines may be more likely to *solve*, novel tasks.

## Introduction

Blue tits opening milk bottles left on the doorstep of English homes is perhaps one of the most well-known animal behaviors, and widely considered an example of animals ‘innovating’ (Laland and Reader 2010). Innovation generates broad interest because it is a critical element of our self-image as humans, and, like ‘intelligence’, seen as a key factor in our evolutionary success (Shettleworth 2012; Reader et al. 2016). Also because of this, and also like intelligence, there are thus widely varying and hotly debated definitions of what actually counts as innovation; but to quote Reader et al., “definitional arguments can provide more heat than light” (Reader et al. 2016). Innovation can refer to using a novel method to solve a problem, or solving a novel problem with an old method (where the blue tit example, like the famous wire-bending of Betty the crow, is more similar to the latter, (Brown 2022). Such behavioral flexibility can particularly improve fitness in changing environments or during biological invasions (Sol et al. 2016; Magory Cohen et al. 2020), with wide ranging consequences from resistance to extinction (Ducatez et al. 2019) to maintenance of behavioral variation in populations (Duckworth and Badyaev 2007).

For an act of innovation to occur, an animal has to encounter an opportunity to innovate, such as a novel potential food source (as the milk bottles). The chance of such encounters may vary with environments (e.g. ecological changes) and individual traits (e.g. movement rates). The animal then has to perceive the opportunity as such and interact with it - whether this is detecting a novel flower or investigating all objects with certain properties. Lastly, a successful innovation implies that the animal ‘solved’ the problem, e.g. employed a novel motor routine to achieve its goal. Typically it is also assumed that some kind of learning or insight occurred either before or after the success occurred (Ramsey et al. 2007; Loukola et al. 2017; Santana and Garcia-Mijares 2022; Brown 2022), so that the innovated activity can be repeated. The environments or traits promoting encounter rates may not be identical to the ones promoting curiosity or engagement with novel stimuli; and those in turn may or may not relate to the ones leading to ‘success’ (Cole et al. 2011; Brosnan and Hopper 2014; Tebbich et al. 2016; Maák et al. 2020). In laboratory studies in particular, it is therefore more informative to identify which part of the innovation process is being studied.

In addition to defining what constitutes an ‘innovation’ and which step on the path to innovations is being studied, investigators have to confront the problem of determining what can be considered ‘novelty’ (Heger et al. 2019; Santana and Garcia-Mijares 2022; Brown 2022). Defining ‘novelty’ is not as straightforward as it seems: even a naive individual, who has never encountered a particular problem or object before, may be ‘evolutionarily prepared’ for it (Heger et al. 2019). Solving problems commonly encountered by the species is not generally considered ‘innovation’ even if the individual who solved them has not performed the same task before. Moreover, animals readily generalize across a variety of dimensions (Ghirlanda and Enquist 2003; Xu and Plowright 2022), such that different problems that are all new to the individual may still require different degrees of ‘innovation’, if particular problems or solutions resemble those the animal has personally experienced or is evolutionarily prepared for. The famous examples of blue tits and crows both involve motor routines that are employed by those species in other extractive contexts, and hence require more ‘transfer’ of knowledge than ‘innovation’ (Brown 2022). Nonetheless, some species, and perhaps individuals, are clearly more predisposed to innovate, even when the problem to be solved only involves a moderate amount of ‘novelty’ (Alem et al. 2016; Prasher et al. 2019).

Here we are interested in what contributes to this variation, in other words, when is innovation most likely to happen? We use an extractive foraging problem-solving task with objects novel to the individuals encountering them, the typical situation for empirical studies of innovation (Griffin and Guez 2014). Here, lab-raised bumble bees extract sugar solution from artificial flowers. These generalist foragers are at least somewhat evolutionarily prepared for ‘solving’, i.e. extracting nectar from, a variety of complex flowers, making it likely that a number of our subjects will succeed with the task (Mirwan and Kevan 2014; Muth et al. 2015; Alem et al. 2016; Loukola et al. 2017; Barker et al. 2018). We investigate which factors shape the frequency with which individuals succeed in solving these novel problems, as well as which steps towards a successful solution are impacted. Two broad categories of factors have been proposed: the environment, and individual personality traits (Amici et al. 2019; Kikuchi 2024; Lermite et al. 2017; Magory Cohen et al. 2020). We focus in particular on environmental complexity.

Seemingly contradictory effects of environmental complexity on animal cognition have been proposed. Visual and spatial complexity of an animal’s surroundings could interfere with cognitive processes needed to solve particular problems (Dunbar 1992; Lefebvre et al. 1997); in particular, stimulus detection is likely to be impeded by distractors (irrelevant visual features; (Chittka et al. 2003; Skorupski et al. 2006; Ohashi and Thomson 2013)). In addition, complex environments may occupy cognitive resources that then become unavailable to solve novel problems (Clark and Dukas 2003). On the other hand, complex environments may aid innovation by decreasing routine formation and prompting animals to remain flexible and pay attention to a larger variety of stimuli (Shumway 2008; Wang et al. 2017). For instance, pigs raised in enriched environments are more likely to notice changes in cues during a task (Bolhuis et al. 2004), and variability in food resources experienced tends to increase behavioral flexibility (Sol et al. 2005). It may thus be difficult to extract generalities on how environments, and particularly environmental complexity, affect innovation.

By comparison, the role of individual differences - sometimes referred to as ‘personalities’, ‘coping styles’, ‘tempo’, or ‘pace-of-life’ - in solving novel problems seems obvious (Biondi et al. 2010; Cole et al. 2011; Overington et al. 2011; Prasher et al. 2019). Foundational work on individual differences since the 1970s has included an exploration-vs.-exploitation dimension (Wilson et al. 1994; Koolhaas et al. 1999; Biro and Stamps 2010; Hills et al. 2015; Toscano et al. 2016; Lemanski et al. 2019). ‘Exploratory’ individuals are thought to show greater variation in behavior and movement and attend more to novelty, thus predisposing them to be ‘information producers’ (Toscano et al. 2016; Lemanski et al. 2019) and thus also, perhaps, to be innovators (Overington et al. 2011; Tremmel and Müller 2013; Udino et al. 2017; Prasher et al. 2019; Collado et al. 2021). After the initial enthusiasm for finding non-human ‘personalities’ gave way to a wealth of detailed empirical studies, however, it became clear that a number of traits such as ‘exploration’, ‘boldness’, ‘responsiveness’, ‘neophobia’, ‘cognitive ability’, ‘flexibility’, ‘tempo’, and ‘activity level’ were either defined inconsistently (for the same term) across studies and species, or (for different terms) measured operationally in ways that are so similar that it is not surprising that traits are found to be correlated. It is thus difficult to know what to make of the fact that many studies have found inconsistency across species or contexts in which traits form a unified ‘syndrome’ that contributes to innovation. For example, in solitary bees, exploration, along with shyness and overall activity level, seems to predict innovative problem-solving, but independently of learning ability (Collado et al. 2021). Overall, the relationship between tendency to explore novel environments and innovation can be positive (Collado et al. 2020; Overington et al. 2011; Prasher et al. 2019; Quinn et al. 2016), negative (Lermite et al. 2017), or neutral (Biondi et al. 2010; Cole et al. 2011). Similarly, neophobia, measured as latency to approach a familiar feeding place when a novel object is placed nearby, has been negatively associated with innovation (Biondi et al. 2010; Magory Cohen et al. 2020; Overington et al. 2011), but this is not always the case (Lermite et al. 2017).

We aimed to test several specific hypotheses regarding the effect of environment and individual traits on novel problem solving. We used bumble bee (*Bombus impatiens*) foragers. Bumble bees are generalist foragers that can be completely raised in the laboratory; we could thus assess their ability to solve novel problems, i.e. access reward on (artificial) flowers they were both naive to as individuals and which were at least somewhat dissimilar to any flowers they may have been evolutionarily prepared for. In addition, since foraging took place in an arena, we could also manipulate the environment in which the bees solved these problems.

First, we distinguish between the following three hypotheses regarding environmental effects: (1) If complex environments decrease overall innovation by reducing the detection rate of novel resources, then bees in more complex habitats may not necessarily differ in handling time once they land on novel flowers, but will take longer to find them amid distracting stimuli. (2) If decreased innovation in complex environments is due to individuals suffering from more general cognitive interference, then bees in these environments should take longer to solve a novel flower, regardless of search time. (3) If, on the other hand, complex environments aid innovative ability because of reduced routine formation, bees adapted to a complex environment should be more likely to solve novel problems than those used to simple environments.

Second, we examined several measures of foraging behavior in the bees prior to being confronted with the novel problems for their relationship with innovation. For each of these measures, we examined whether they were (a) affected by the foraging environment and (b) whether they predicted the probability of solving or the time to solve the novel problems. We specifically examined the (i) first handling time on any of our artificial flowers (Muth et al. 2015; Barker et al. 2018), (ii) the search time in the first innovation trials (Chittka et al. 2003), (iii) the overall travel time per trip prior to innovation trials, (iv) ‘responsiveness’, a measure used in the literature that may indicate curiosity or distractability (Benus et al. 1987; Mathot et al. 2012; Wolf et al. 2008), (v) routine formation (Ohashi and Thomson 2013), and (vi) explorative tendency (Hills et al. 2015; Collado et al. 2021). While we name these traits for ease of reference and consistent with prior literature, it is important to keep in mind that they are based on simple measurements of behaviors or behavioral outcomes; what mental process they exactly reflect often remains uncertain, in our study and other similar studies. Nonetheless, intuition, prior studies, and our results all suggest that, in bees, the process of flying and landing on flowers (important for ii, iii, iv, and vi) is distinct from the process of finding and accessing nectar on a flower after landing (i); and visiting known flowers in a specific sequence known as ‘trapline’ (v), is a well-described specific behavior in bees. Our primary goal was to identify whether any behaviors measured prior to the actual process of accessing reward on a novel flower (what we term ‘innovation’ here) would predict the outcome, implying that innovation is affected by traits of the bee and not just a chance occurrence.

## Methods

### Bee set up and care

Four colonies of *Bombus impatiens* were procured (Koppert Biological, USA) and kept in wooden nest boxes measuring 39 cm x 10.5 cm x 23 cm where they were maintained on a substrate of Arm & Hammer Feline Pine Original Cat Litter. Nest boxes were covered by clear acrylic plastic lids. The boxes were connected to a wooden foraging arena (75 cm x 60 cm x 40 cm) by clear acrylic tubing with dividers allowing us to control access to the arena (see Jandt and Dornhaus 2011 for details). On non-experiment days we fed the bees a 25% (by volume) sucrose in water solution in a transparent glass/plexiglass gravity feeder placed in the center of the foraging arena, raised off the surface by 8 cm on a plastic cylinder to motivate the bees to fly. The colony was allowed free access to the arena except during the training and experimental phases. We provided ground honeybee pollen (Koppert Biological) directly onto the nest once a day.

### Flowers

Artificial flowers were created using colored sheets of craft foam. In the center of each flower, we placed a 100 µL microcentrifuge tube to hold a sucrose reward. We glued metal wire to the bottom of the flowers and placed the flowers in plastic vials or rubber stoppers such that the ‘corolla’ was held horizontally. The four flowers used in Trials 1-4 (hereafter called standard flowers) were yellow, flat, and circular with a diameter of the corolla of 3.8 cm (Figure 1A). In Trial 5 we used an additional three blue flowers that were identical in size and morphology to the standard yellow flowers. In Trials 6-8b, we used four types of novel flower morphologies (Bumpy, Folded, Cap1, and Cap2; Figure 1). All of these novel flowers were also yellow (Figure 1). The novel flower used in Trial 6 had small alternating folds of foam glued to its flat circular surface. The novel flower used in Trial 7 was made of a rectangular piece of foam folded into an acute angle to create a ‘V’ shape when viewed from the side (Figure 1B). The flowers used in Trial 8a-b was the same as those used in Trials 1-5, except that the lid of the microcentrifuge tube was used to prevent access to the sucrose solution. The lid was placed either upside down on top of the tube so that the bee had to push the cap aside (Trial 8a) or partially inserted in the tube so that the bee had to pry open the cap (Trial 8b). Bees did not leave the arena between 8a and 8b, instead flower 8b was swapped in once the bee left the flower 8a and so we consider it to be a single trial with two parts. These novel flowers challenged bees to use a variety of cognitive and motor abilities (from tasks such as navigating between bumps, walking on vertical incline, manipulating the cap) and are within the range of task difficulty that bumble bees have been shown to master (Alem et al. 2016; Loukola et al. 2017).

**Figure 1.**
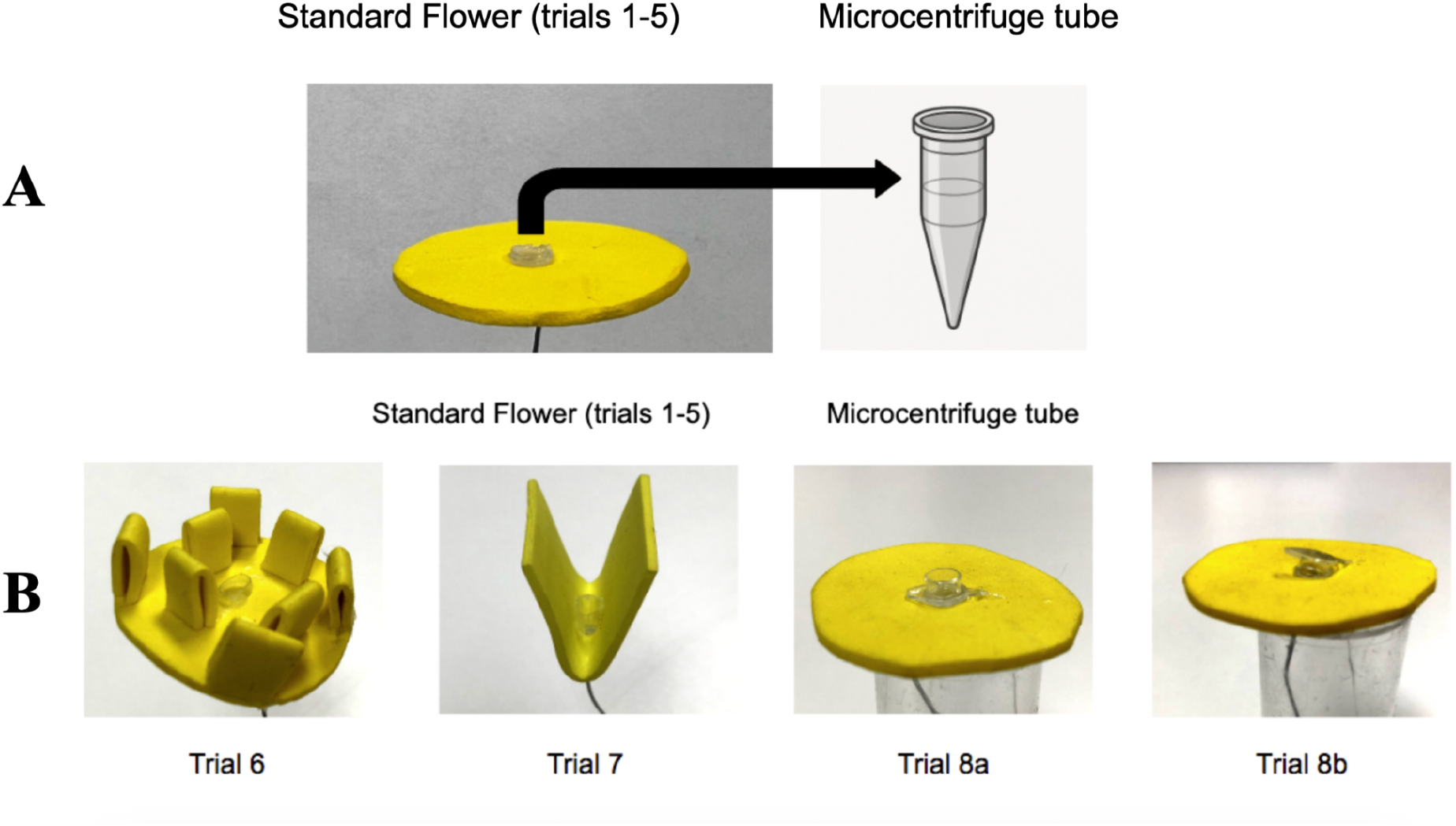
(A) Standard flower used in trials 1-5 with a schematic of the microcentrifuge tube that was mounted in the center of all flowers. (B) Flowers used in the novel flower trials. Trial 6 used the Bumpy flower, trial 7 used the Folded flower, trial 8a used the Cap1 flower (cap, placed upside down, needs to be pushed off to access nectar) and in trial 8b, with the same flower type, the cap needed to be pried off to access nectar: we call this treatment Cap2.

### Pre-training

All individual workers in each colony were marked on the thorax with a colour- and number-coded plastic tag (Queen Numbers Set, Betterbee, Greenwich, NY). To familiarize bees with the general foraging task, we removed the gravity feeder and added 10 standard yellow flowers to the foraging arena. Each flower was filled with 25µL of 50% (by volume) sucrose in water solution to motivate bees to forage from the flowers. We manually replenished the flowers as they became depleted by the free-foraging bees (at the discretion of the observer). To begin a training session, all bees were returned to the nest from the foraging arena and flowers were spaced evenly throughout the arena. We allowed only labeled foragers access to the arena. Bees that did not show interest in foraging from the flowers within 10 minutes of entering the arena were removed and placed back in the nest. When an individual was able to collect sucrose solution, deposit it at the nest, and return to the arena within 5 minutes we removed other bees from the foraging arena and began the experimental trials with that bee.

### Experiment

During trials, only the bee being tested was allowed in the arena. Bees were pseudo-randomly assigned to one of two different environmental complexity treatments, which then applied for all experimental trials (Figure 2). Each bee experienced eight experimental trials that are described below, returning to the nest after each trial. We refilled flowers after each trial and washed them with isopropyl alcohol and water after a bee completed all trials. We refer to trials 6, 7, 8a, and 8b as test trials because in these trials bees were presented with novel rewarding flowers (where ‘innovation’ might be needed to access reward). In the simple environment treatment, the arena was uncluttered, with only the experimental flower(s) present. In the complex environment treatment, the arena was cluttered with artificial plastic ivy (a kind of garland of stalks and leaves) and 10 non-rewarding distractors (2.54 cm yellow craft foam squares), both of which were spread evenly on the arena floor (Figure 2B). Flower arrangement was identical across treatments.

**Figure 2.**
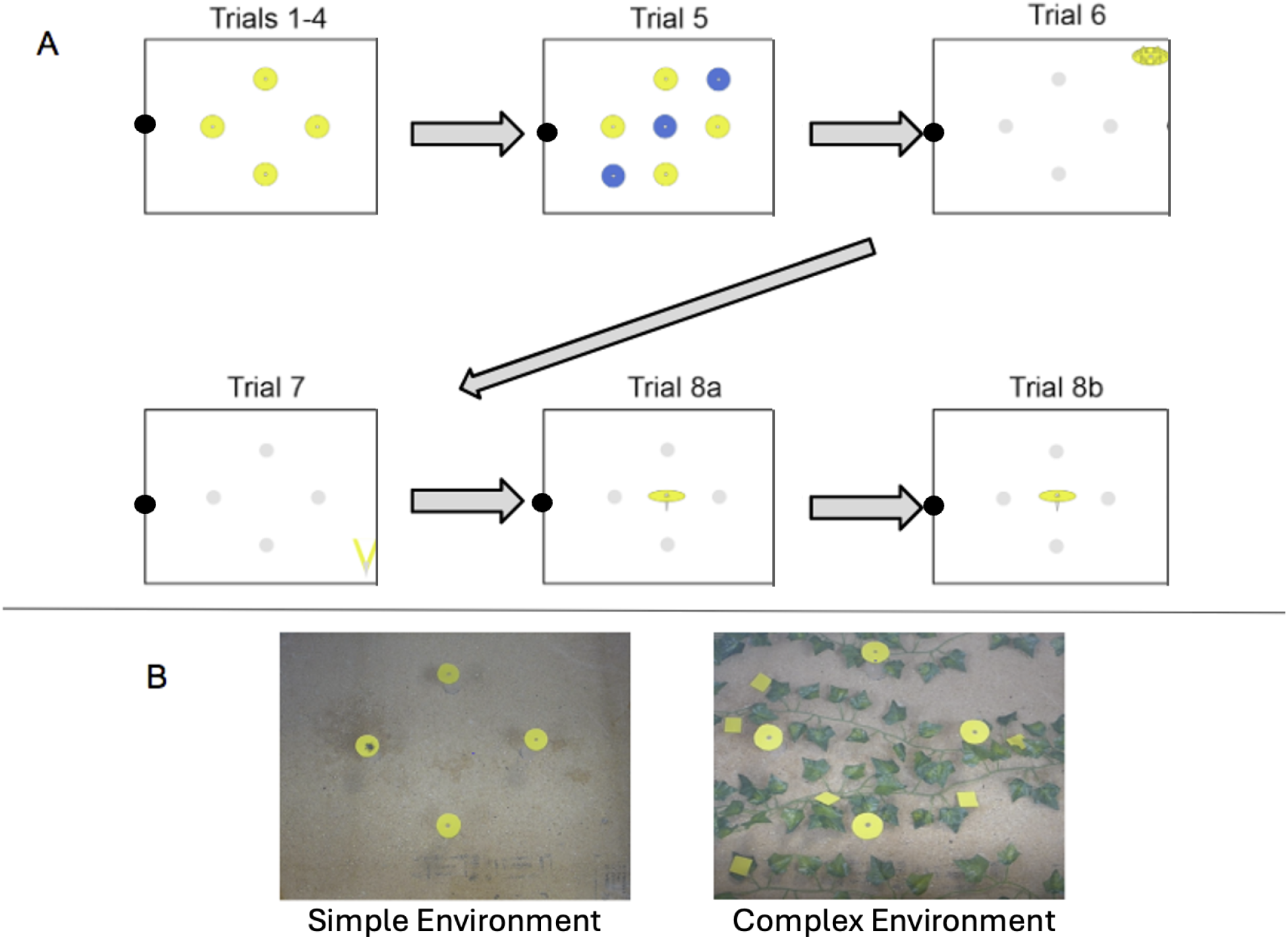
(A) Schematic showing the order (grey arrows) and flower arrangement in each trial viewed from above the arena. The black circle shows the entrance to the arena. Blue and yellow circles represent the standard flowers and the non-circular yellow shapes represent the flowers used in Trials 6, 7, 8a, and 8b respectively. Gray circles represent overturned plastic vials (which previously held the yellow flowers). Trial 8b occurred immediately after Trial 8a, without the bee returning to the nest. (B) Example images of the foraging arena in simple (left) and complex (right) treatments.

Each trial began when the bee entered the arena. If bees were unable to locate or access the reward in the flower(s) within 10 minutes of the beginning of the trial, the trial was recorded as incomplete. After an incomplete trial we waited for the bee to leave the arena. The bee was then allowed to proceed to the next ‘trial’, i.e. task as per Fig. 2, but no outcome was scored for the trial that was incomplete. No bee was allowed to repeat any trial. When a bee had completed all trials it was removed from the colony and frozen.

Specific flower arrangements for the 8 trials:

**Trials 1 – 4:** These trials were identical and established a baseline to compare to trial 5 for our measure of responsiveness. Four standard flowers were placed in a diamond shape with 23 cm between vertices (Figure 2). Each contained 25µL of 50% (1:1 by volume) sucrose water (this concentration was used throughout the study). This total volume was enough for most bees to immediately return to the nest after foraging from the fourth flower. Flowers were exchanged with new, identical, filled flowers to minimize scent marks and resulting effects on bee choices. **Trial 5:** In addition to the flowers present in Trials 1 – 4, three non-rewarding blue flowers were added. They were arranged diagonally between the yellow flowers (Figure 2). The blue flowers served as a novel stimulus and were used to measure the bees’ responsiveness and exploration tendencies.

**Trials 6 – 8:** all flowers from previous trials were removed. The vials that held the yellow flowers were flipped upside down and left in the arena (but did not now contain any yellow stimuli). To decrease location bias from prior foraging, the flowers for trials 6-8 were placed in one of the four corners of the arena, areas which had not previously held a rewarded flower as per Fig. 2. There was only one of each novel flower, containing 25 µL sucrose solution, presented in each trial, as follows:

**Trial 6:** One novel Bumpy flower containing 25 µL sucrose solution was placed in one corner of the arena with 100 µL sucrose solution.

**Trial 7:** One novel Folded flower containing 25 µL sucrose solution was placed in a different corner of the arena with 100 µL sucrose solution.

**Trial 8a:** Flowers in this trial were placed in the center of the arena. The cap of the centrifuge tube was placed upside down on the tube, requiring the bee to push it off of the opening to access the reward. It contained 25 µL sucrose solution so that the bee was still motivated to forage in trial 8b. After the bee completed this task (or 10 minutes passed), trial 8b was initiated while the bee was still in the arena.

**Trial 8b:** Before the bee left the arena, we quickly replaced the flower with an identical one with 25 µL sucrose solution that had the cap set into the tube gently (the correct way – down). This required the bees to use its body to remove the cap off of the tube to access the reward.

### Video Analysis

All trials were recorded from above the clear acrylic cover of the arena using either a Panasonic Lumix LX-3 camera or a GoPro Hero Session 4 camera. The videos were viewed on a color monitor using VLC Media Player software on a PC.

The total duration of each trial was defined as the time from the moment the bee appeared at the arena entrance to the moment the bee disappeared upon exiting the arena. We subdivided each trial into two main components: travel time and handling time. Travel time was defined as the time a bee spent moving between flowers from the time the bee entered the arena until it landed on the last rewarded flower, while handling time was the time spent while on any of the rewarded flowers (i.e. not flying, and touching the flower with at least 2 legs). In the non-innovation trials (trial 1-5), handling time includes time spent drinking at the flower. In the innovation trials, we aimed to measure the time it took to access the nectar, so handling time in trials 6-8 only reflects time from landing on a flower (at least 2 legs on flower) until the bee began drinking (insertion of proboscis into well). If the bee did not reach the reward, we quantified the total time on the flower (until the bee left) as ‘giving up time’.

### Data analysis

All calculations and analyses were performed in R version 4.5.0 (R Core Team 2024) using RStudio (RStudio Team 2025). We used the lme4 package to run regressions (Bates et al. 2015).

Our main aim was to quantify the impact of a simple or complex environment on innovation, but we also analyzed the impact of several individual traits on innovation, and the impact of the environments on these individual traits, to explore how both environment and traits may affect innovation. We quantified 6 behavioral outcomes that could be seen as reflecting individual traits: (1) handling time on the first flower in trial 1; (2) search time (average time from entering the arena to landing on the flower in trials 6&7); (3) travel time (total travel across all flowers in trials 2-4), a measure we interpret as foraging speed; (4) routine formation; (5) responsiveness; and (6) exploration. We defined routine formation as the degree to which bees stuck to a fixed trapline, i.e., a repeatable sequence in which they visited individual flowers, a foraging pattern common in bees (Lihoreau et al. 2011; Ohashi and Thomson 2009). Routine formation was quantified using the Sequence Repetition Index (SRI), which was derived from the Levenshtein distance (Levenshtein 1966) between foraging sequences of trips 1-4 using the stringdist package in R (Van Der Loo 2014). Responsiveness was calculated as the proportional increase in travel time between the average of trips 2-4 and trip 5 (where the blue flowers were introduced). We averaged trips 2-4 in order to even out any variability in travel time for each bee and we excluded the first trial as the travel (and search) time would naturally be longer their first time in the arena with this configuration of flowers. We assume here that bees that spent more time traveling in trial 5 as compared to trial 4 were displaying some sort of interest in or distractibility by the blue flowers, compared to bees that may have completely ignored the blue flowers, just foraging on the yellow flowers as before (and thus keeping their travel time the same). In addition, we categorized bees as having explored blue flowers if they ever landed on the blue flowers; we termed this binary category ‘Exploration’.

We used binomial logistic regression to assess the effect of environment and flower type on the probability of landing on (Table S1) and solving (Table S2) the novel flower. We also tested whether individual traits predicted the proportion of trials in which a bee landed on or solved the novel flower using linear regression (model: proportion landed or solved ∼ trait score).

Waiting times are usually exponentially distributed (e.g. with normally distributed probability of success per time), and indeed the distribution of time to solve the task (our measure of innovation) resembled an exponential distribution more than a normal distribution (Figure S1). We therefore log-transformed all time values (i.e. used log(time) as the response variable in models). To assess the effect of environmental complexity on innovation, we fit a linear model with time to solve as a function of the environment treatment, trial, and their interaction (log(time) ∼ env * trial, where trial also reflects the flower type bees are ‘solving’). To test whether individual traits of bees affected this time to solve the task, we used individual models of the form log(time) ∼ trait * trial, i.e. with both the trait in question and the trial as well as their interaction as factors. We generally give full results tables for models in the Supplemental Materials.

To determine whether the environment had an effect on individual bee behaviors other than innovation, we performed individual Mann-Whitney-U-Tests comparing the median trait in bees in the simple against that of bees in the complex environment.

To separately determine whether we would detect significant within-bee repeatability of time to solve, we first fit a linear mixed-effects model with bee ID as a random intercept and trial as a fixed effect, using the lme4 package in R. We then compared this model to a reduced model excluding the random effect, using a likelihood ratio test (LRT) to evaluate whether including individual identity improved model fit. Second, we estimated the repeatability (intra-class correlation coefficient, ICC) of handling time using the rptR package, which partitions variance into between- and within-individual components.

## Results

### Landing and solving

We observed 35 bees in total (18 in the simple, 17 in the complex environment) across the 9 trials (of which 4 trials were testing ‘innovation’). Twenty-nine bees had an observation for each of the four novel flower trials (meaning they at least landed on each flower in trials 6, 7, 8a, and 8b) while 6 bees were missing one observation (meaning they did not even land on at least one trial flower before the time ran out or they returned to the hive). Bees were overall able to solve the task, i.e. drink sugar solution from a novel flower, in 89% of trials in which they attempted it (landed) across all novel flower types (117/132). We find that bees in complex environments were significantly less likely to land on the novel flower compared to those in simple environments (z = -1.98, p = 0.048; Figure 3, Table S1), but environment did not significantly affect whether bees solved the flower after landing (z = -0.64, p = 0.53; Figure 3, Table S2).

**Figure 3.**
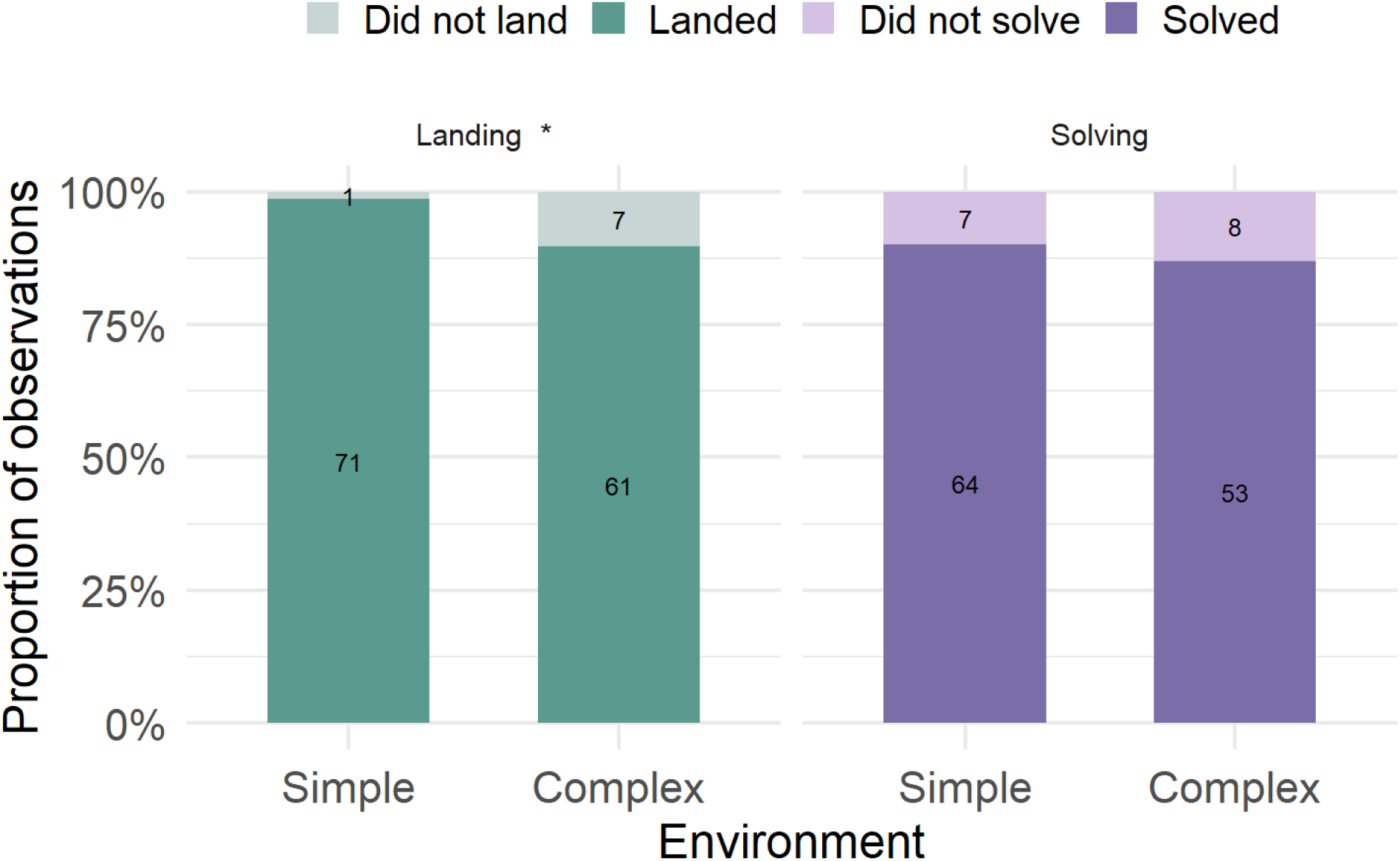
Proportion of observations of landings on a novel flower (left) and instances of solving the novel flower task once landed (right), shown by environment treatment. Numbers within bars represent the number of observations for that category; individual bees had up to four observations (across different trials). In the binomial linear models, bees in the simple environment were found to be significantly more likely to land on the novel flower compared to those in the complex environment (p = 0.048), but the environment did not significantly affect the probability of finding the reward among bees that landed (p = 0.525). Landing model: landed (y/n) ∼ environment + trial. Solving model: solved (y/n) ∼ environment + trial.

It seems that the order of novel flower types we chose reflected their complexity for bees: The majority of failures to solve were observed in the Cap2 flower (7 bees out of 35), followed by Cap1 (4 bees), Folded (3 bees) and then Bumpy (1 bee). In addition, Cap1 and Cap2 differed significantly in our main measure of ‘innovation’, i.e. the time to reach reward in the novel flowers once landed, from ‘Bumpy’ and ‘Folded’ (relevant pairwise comparisons p<0.0001; Fig. 4, Table S3). Since only a few bees gave up (abandoned flowers after landing without extracting reward), we can say little about what determines the time to giving up, but this also seemed to increase with flower complexity (Figure S2) and be independent of environment (Table S4).

**Figure 4.**
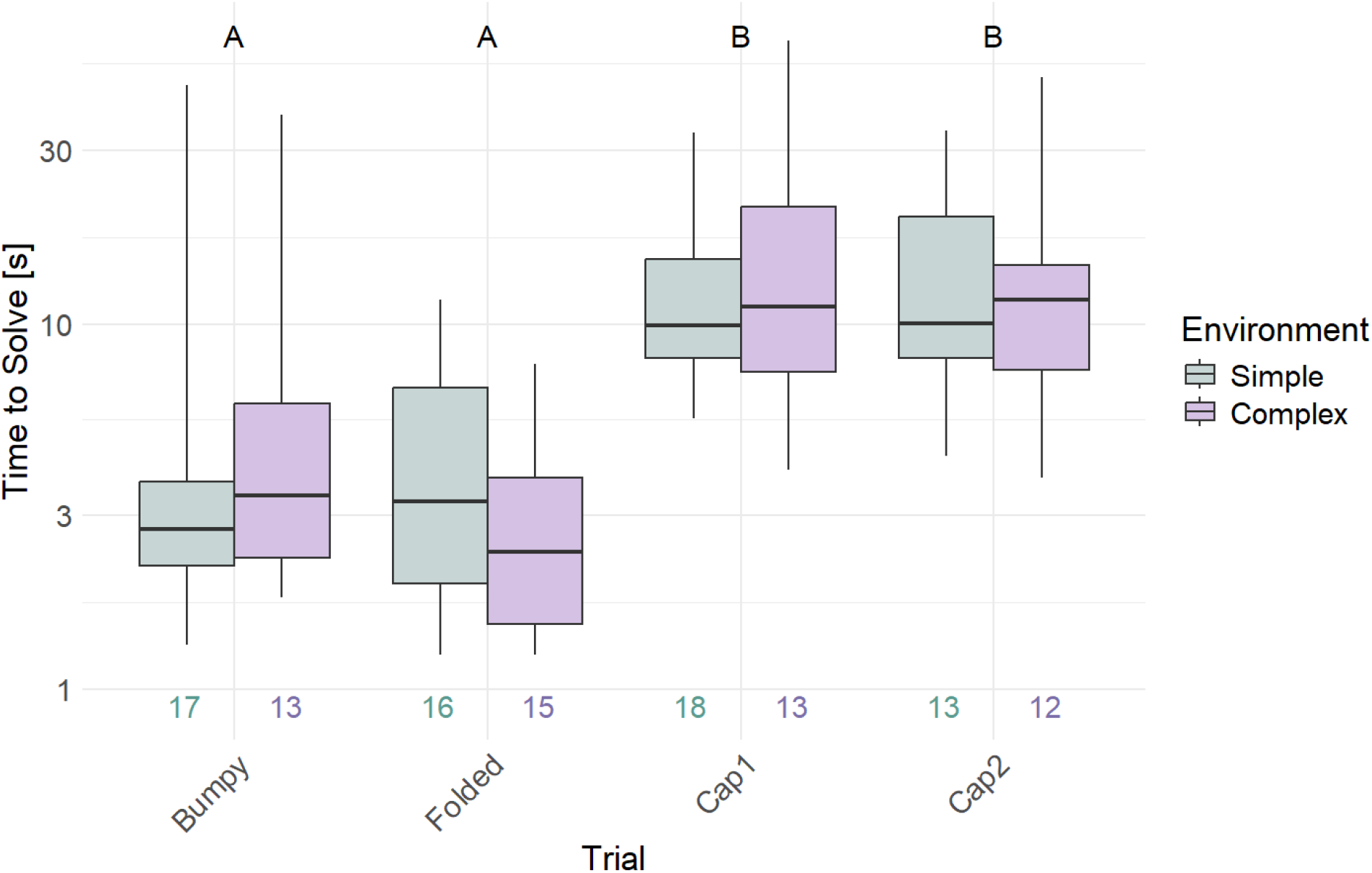
The time it took bees to solve the novel flower, i.e. extract reward once landed (y-axis), separated by trial (x-axis) (sample sizes are number of bees). Bees were tested in either a simple or complex environment (colors). Environment did not significantly affect solving time (p = 0.52), but flower type did (p < 0.001). Letters A and B indicate statistically significant differences in handling times between flower types. Model: log(time to solve) ∼ env * trial.

With the exception of search time, none of our individual traits (first handling time, travel time, routine formation, responsiveness, or exploration) significantly predicted the probability to land (p-values 0.27, 0.75, 0.24, 0.93, 0.26, respectively) or ‘solve’ novel flowers (p-values 0.11, 0.09, 0.16, 0.35, 0.27 respectively). Search time (averaged over the Bumpy and Folded flower trials, which were the ones where flowers were placed in novel locations) significantly predicted both landing and solving (p < 0.0001 for search time on landing, and p = 0.001 for search time on solving; Figure 5). Bees who searched longer (time from entering arena to landing) overall landed less often and solved novel flowers less often (Figure S3).

**Figure 5.**
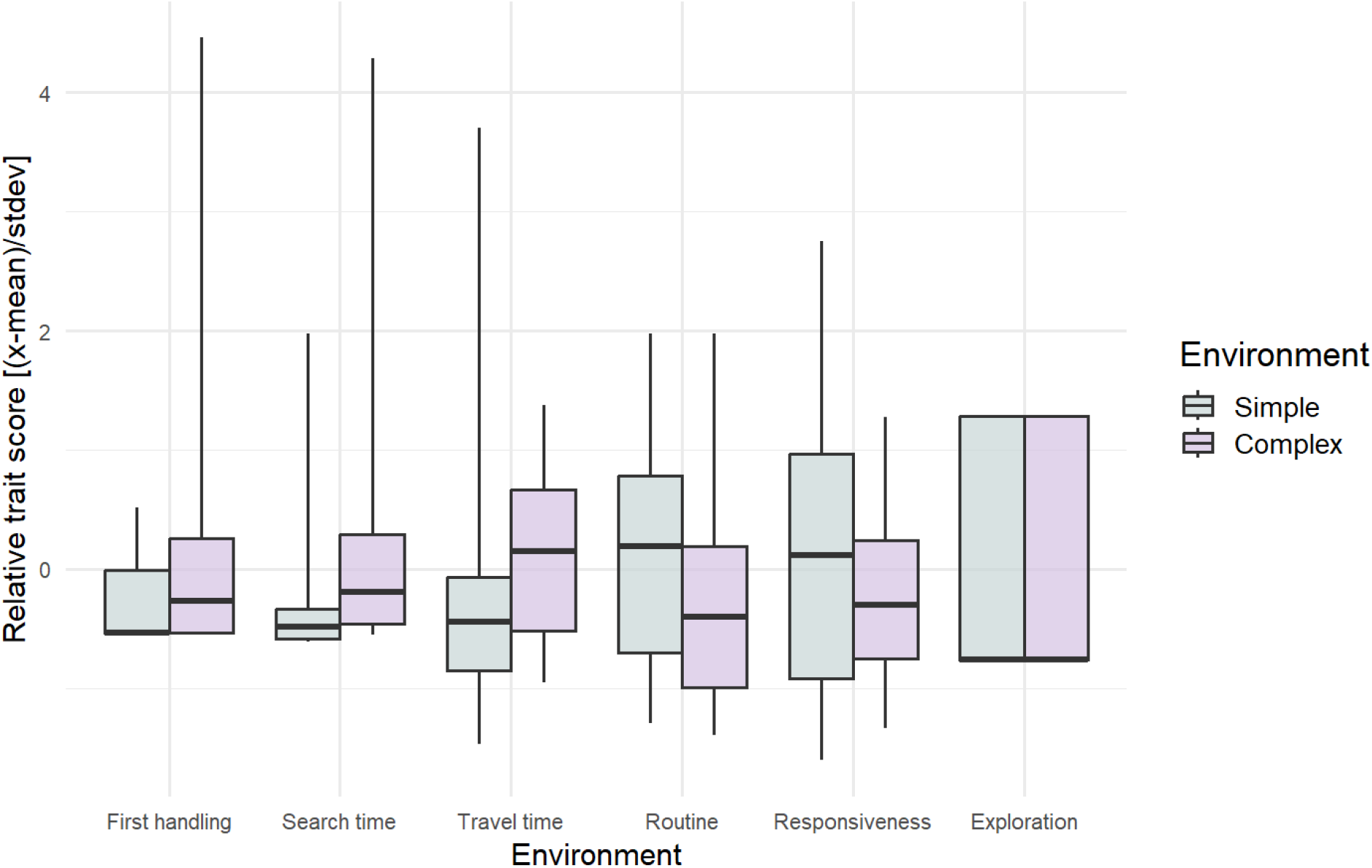
Individual bee behaviors or traits in relation to the environment the bees were tested in (across all 9 trials). Search time was longer in the complex environment; other traits were not affected by environment (Mann-Whitney-U-Tests, for the 6 traits p = 0.055, 0.02, 0.09, 0.10, 0.66, 0.84). The y-axis shows the relative trait score = (actual score - mean for that trait)/(standard deviation of scores for that trait).

**Figure 6.**
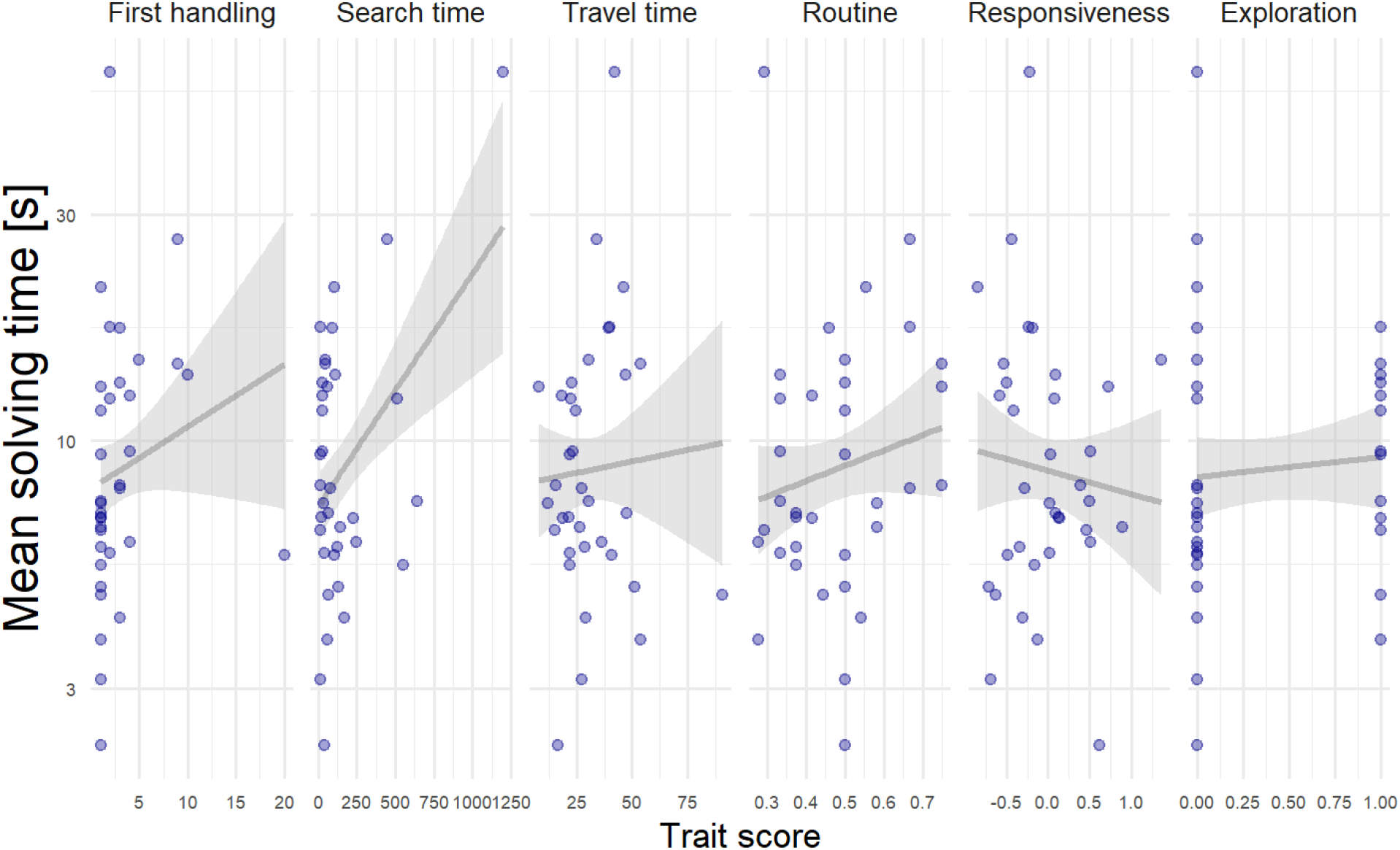
Relationship between average solving time (across all innovation trials, 6-8b) and individual traits (measured in earlier trials, except for search time). ‘First handling time’ (handling time on very first flower, in trial 1); Search time (in trials 6 and 7); Travel time (across trials 2, 3, and 4); Routine formation (SRI) (in trials 2-5); Responsiveness (increase in foraging time upon addition of new stimuli in trial 5), and Exploration (landing on blue flowers in trial 5). Each point represents a bee’s mean solving time across the 4 innovation trials (y-axis, logarithmic) and the bee’s trait score (x-axes). Lines show linear regression fits with 95% confidence intervals; only travel time and routine formation significantly predicted solving (First handling: p = 0.06, Search time: p = 0.34, Travel time p = 0.02, Routine formation p = 0.02, Responsiveness p = 0.13, Exploration p = 0.31). All models: log(solving time) ∼ trial * trait.

### Effects of simple vs complex environment on bee behavior

The simple or complex environment in our trials (Figure 2) only affected search time significantly, with bees in the complex environment searching longer (Mann-Whitney-U-Test, W = 75, p = 0.017; Figure 5). The first handling time (p = 0.055), travel time (p = 0.09), routine formation (p = 0.10), responsiveness (p = 0.66), and exploration (p = 0.84) were not significantly affected (Mann-Whitney-U-Tests, all n(simple) = 18 and n(complex) = 17).

### What predicts innovation?

We sought to compare the effect of environment with the effect of an individual bee’s traits on innovation, here measured as time to reaching reward on novel flowers (once landed on them). Environmental complexity (i.e., whether or not distractors were added) did not significantly affect solving time (t = 0.65, SE = 0.28, p = 0.52; Fig. 4, Table S3).

In contrast, some of the individual bees’ behaviors prior to solving the novel flowers did predict ‘innovation’, i.e. the time to reach the reward on these novel flowers. In particular, travel time, the time bees spent flying between flowers in trials 2, 3, and 4 (i.e. on the plain yellow flower set), predicted innovation, with longer travel time predicting longer solving time (t = 2.39, SE = 0.26, p = 0.018), perhaps indicating overall differences in foraging tempo among bees. In addition, SRI (sequence repetition index), a measure of routine formation, was significantly positively associated with solving time (t = 2.46, SE = 0.97, p = 0.016; Table S5, Figure 5), indicating that bees with more repetitive foraging sequences in earlier trials took longer to solve novel tasks in the innovation trials. The relationship of responsiveness, defined as the increase in time spent in the arena following the addition of novel stimuli, with solving time was not as clear: while there was a negative trend, responsiveness as a main factor did not have a significant relationship with solving time (t = -1.55, SE = 0.25, p = 0.13), but interacted significantly with trial to affect solving time (resp x Folded t = 2.01, SE = 0.35, p = 0.047, Table S7). Handling time on the very first (plain) flower of the first trial just failed to significantly predict solving time (t = 1.91, SE = 0.16, p = 0.059, Table S6). Search time for novel flowers (t = 0.96, SE = 0.15, p = 0.34) and the exploration trait, whether or not the bee interacted with the novel blue flowers in trial 5, did not significantly predict solving time (t = 1.02, SE = 0.27, p = 0.31).

Overall, a likelihood ratio test comparing a mixed-effects model with BeeID as a random intercept to a fixed-effect-only model found no significant improvement in model fit when including individual identity (LRT: χ^2^ = 0.70, df = 1, p = 0.402). Consistent with this, repeatability analysis found no evidence of individual differences in solving (handling) time.

## Discussion

A major goal of our study was to test differing hypotheses from the literature on whether and in what direction environmental complexity would affect innovation. We found that our manipulation of environmental complexity did not significantly influence the probability of solving a novel task nor the time it took once a bee had landed. However, bees in complex environments were significantly less likely to land on the novel flower in the first place, and took longer to search for it when they did. This suggests that environmental complexity may limit the opportunity to innovate rather than the ability to do so once the opportunity presents itself, supporting our first hypothesis but not the second and third: In cluttered environments, perceptual and cognitive load and distraction may interfere with stimulus detection, reducing the likelihood that bees even engage with a novel task or stimulus. However, our results do not support the hypothesis that this leads to a general cognitive load affecting the ability of bees to solve any problem that has been encountered. Instead, this interpretation is consistent with findings that environmental structure can influence the likelihood of encountering novelty (Lermite et al. 2017; Magory Cohen et al. 2020). Our results also do not support the idea that complex environments may prevent routine formation or enhance problem solving as a consequence of higher alertness to novelty.

Our second major goal was to determine which, if any, traits of individual bees themselves affected their ability to innovate. When confronted with a novel problem, bees that previously followed a stricter routine, i.e. a particular sequence or ‘trapline’ in gathering nectar from known resources, took longer to solve it. In addition, bees that previously foraged faster in the sense of having a shorter travel time among flowers, solved the novel problem more quickly. Other aspects of the bees’ behavior, such as previous handling time on a simple flower, responsiveness to a change in the visual environment, search time, or exploration (landing on novel flower types, perhaps equivalent to neophilia), had no clear effects on time to solve the novel problem. Our results are overall consistent with an interpretation that bees who were faster foragers and who were less likely to settle into a routine were faster to solve novel problems.

Overall, we thus conclude that the complexity of the environment may affect innovation primarily through preventing individuals from detecting novel stimuli, and that individuals may differ in their propensity to innovating. We defined ‘innovation’ as successfully extracting a reward from an artificial, novel flower shape; this is consistent with an understanding of innovation as a category of problem-solving. However, the quantification of ‘innovation’ is complicated by the fact that different conceptual and operational definitions have been used in the literature (see Reader et al. 2016; Brown 2022; Bandini and Harrison 2020).

Innovation is likely driven by many of the biases that animals have evolved to respond to stimuli in different contexts, and the coincidence between those biases and ecological opportunity (Greenberg 2003; Griffin 2016). In other words, innovation, like other apparently sophisticated cognitive tasks, is not simply a function of ‘cognitive complexity’ alone. Instead, innate biases and cognitive mechanisms can promote or hinder problem-solving in a variety of ways. For example, in bumble bees, an innate bias for foraging on purple flowers can cause bees to persist and eventually succeed in attempting to gain nectar from them, even if the flowers are of such complex morphologies that the bees are initially unsuccessful (Muth et al. 2015). In other words, a simple innate color bias in this case is a requirement for complex learning. On the other hand, a bias for preferring blue flowers can also prevent honey bees from learning to distinguish between flowers with different shapes (Morawetz et al. 2013). Color bias could lead bees to innovate more frequently on novel purple or blue flowers because the bees persist longer when trying to forage upon them – in other words, obstinate preference may underlie innovation in this case. Similarly, biases not directly related to innovation can affect either the exposure of individuals to innovation opportunities or their success in taking advantage of them. For example, preference for novel stimuli rather than familiar ones (neophilia) could predispose an animal towards encountering novel ecological opportunities, making them more likely to innovate (Greenberg 2003; Greenberg and Mettke-hofmann 2001).

The result that bees who have a less fixed travel route (or ‘trapline’) among flowers in the training trials are more likely to innovate is particularly interesting. This is consistent with literature that individuals with more established routines are less likely to innovate (they are more behaviorally ‘conservative’, (Brosnan and Hopper 2014; Hrubesch et al. 2009). So while traplining is an efficient foraging strategy when resources are stable (Lihoreau et al. 2011; Ohashi and Thomson 2009), it may thus come at the cost of flexibility. In our analysis, solving time was only compared for bees that did land on the novel flower. Slower solving time in bees with stronger traplines therefore implies that routine formation negatively relates to innovation beyond flexibility in the specific movement path. Instead, it suggests that innovation is at least partially affected by the bee’s cognitive state. Our study could not differentiate whether developing a fixed routine in an earlier foraging trip directly prevented bees from innovating, or whether bees that were likely to develop such a routine were generally less prepared to innovate, i.e. had a ‘personality’ that made them less innovative. Whichever is the case, this finding supports the idea that innovation may arise in part from a disruption of behavioral routines (Greenberg 2003; Tebbich et al. 2016).

Overall, our findings suggest that while we did not find strong evidence that environmental complexity supports or prevents innovation, we found that bees that showed slower or more routine foraging behavior tended to take longer to solve novel tasks. Other possible personality traits may also affect innovation, but future work with larger sample sizes or repeated measures could help clarify the timescale across which these behavioral differences among individuals are stable, and the causal factors that cause them to develop. Future studies are also needed to confirm the effects of environmental complexity at different stages in behavioral decision-making. Better understanding how animals approach novel situations could give insight into how they might cope with changing environments or unfamiliar challenges.

## Supporting information

All supplementary figures and tables

## Acknowledgments

We wish to thank other members of the Dornhaus lab for support, and Reuven Dukas, Sabrina McNew, and Dan Papaj for helpful suggestions on the work and the manuscript. DWK and TJP were supported by National Institutes of Health (NIH) training grant 1K12GM000708. DWK was supported by the Wissenschaftskolleg zu Berlin and the German Research Foundation (DFG) as part of the SFB TRR 212 (NC3) – Project number 316099922. AD was supported by NSF, grants no. IOS 3014230 and ABI 3019760.

