## Supplementary material for "Bumble bees that follow a stricter routine innovate less: Foraging behaviors, environmental complexity, and how they relate to novel problem solving": All supplementary figures and tables

### Supplementary Materials

#### Supplementary Figures

###### **Figure S1**: Distribution of solving times (i.e. time from landing to extracting nectar on novel flowers). Dashed line indicates the median.

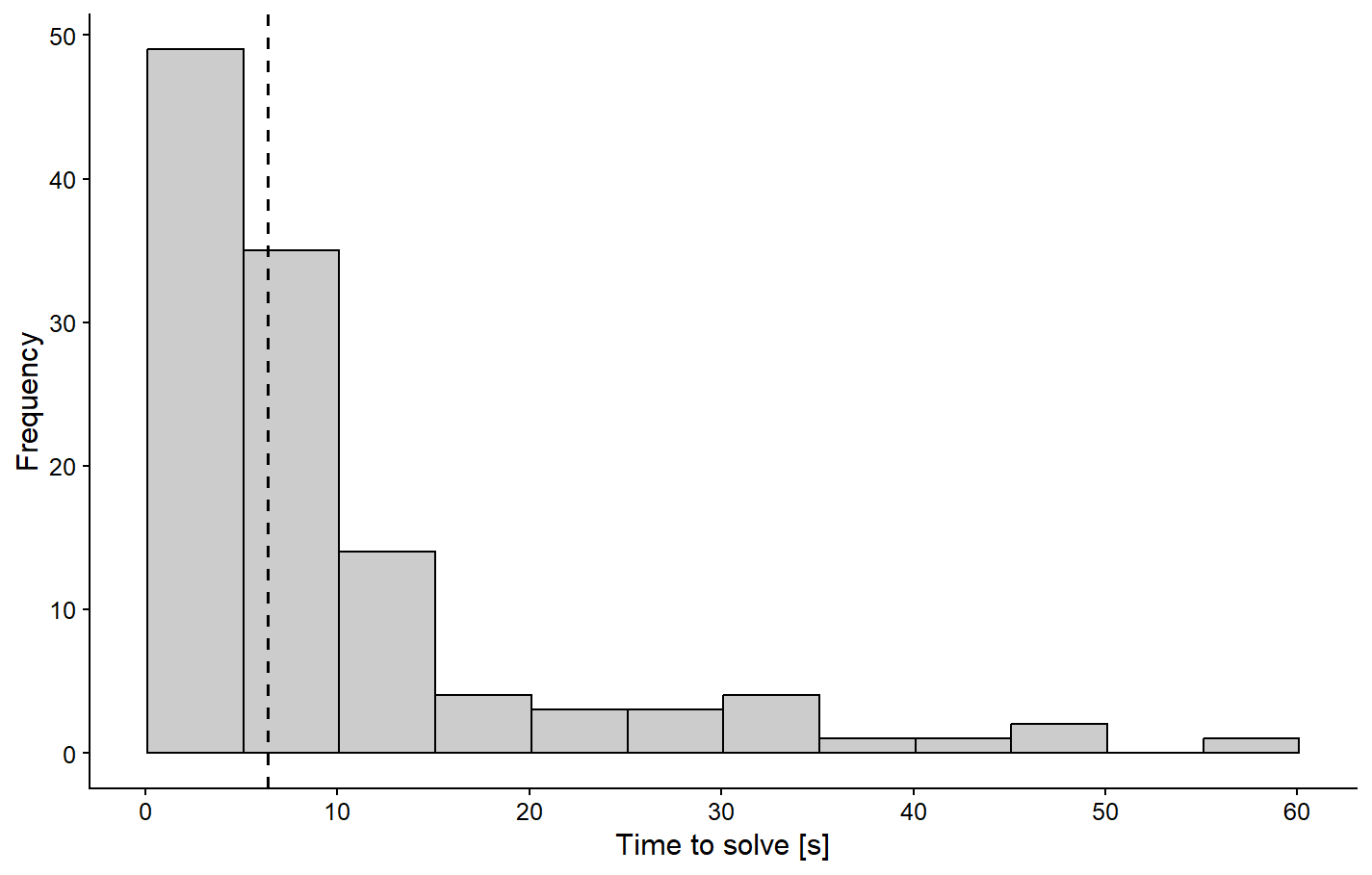

###### **Figure S2**: Time to give up (from landing to leaving flower without extracting reward) in the four innovation trials. Sample sizes (number of bees, shown on x-axis) are small since few bees failed to solve, i.e. extract reward (see also Table S4).

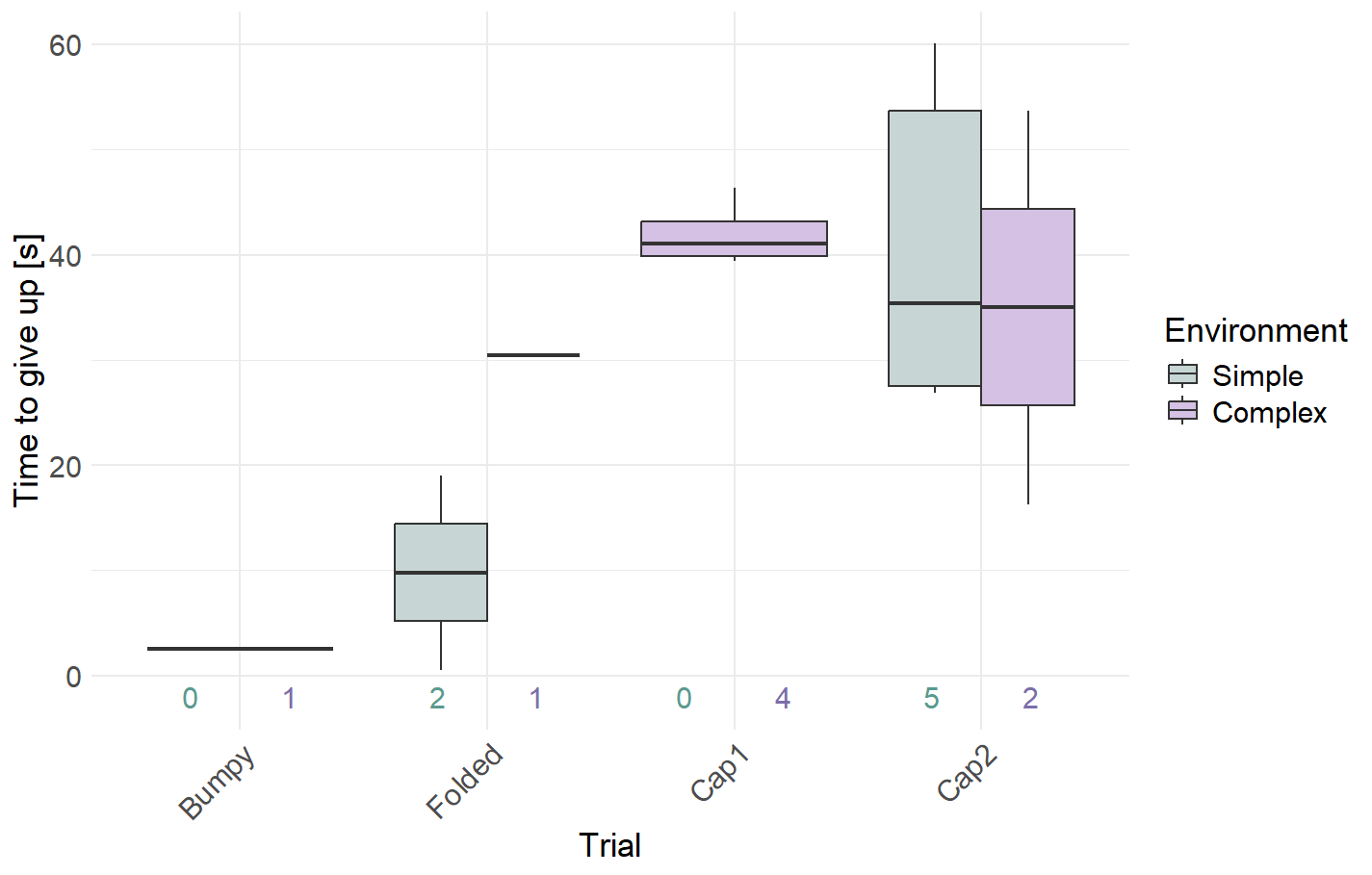

###### **Figure S3**. Relationships between individual traits and behavioral outcomes. Each panel shows the association between a bee’s trait (columns) and the proportion of trials in which bees landed on (top row) or solved (bottom row) a novel flower. Lines show linear model fits. Only search time significantly predicted the probability of landing or solving (Landing: p = 0.273, <0.0001, 0.754, 0.245, 0.93, 0.26, respectively; Solving: p = 0.11, 0.001, 0.10, 0.16, 0.27, respectively). All models: land or solve ~ trait.

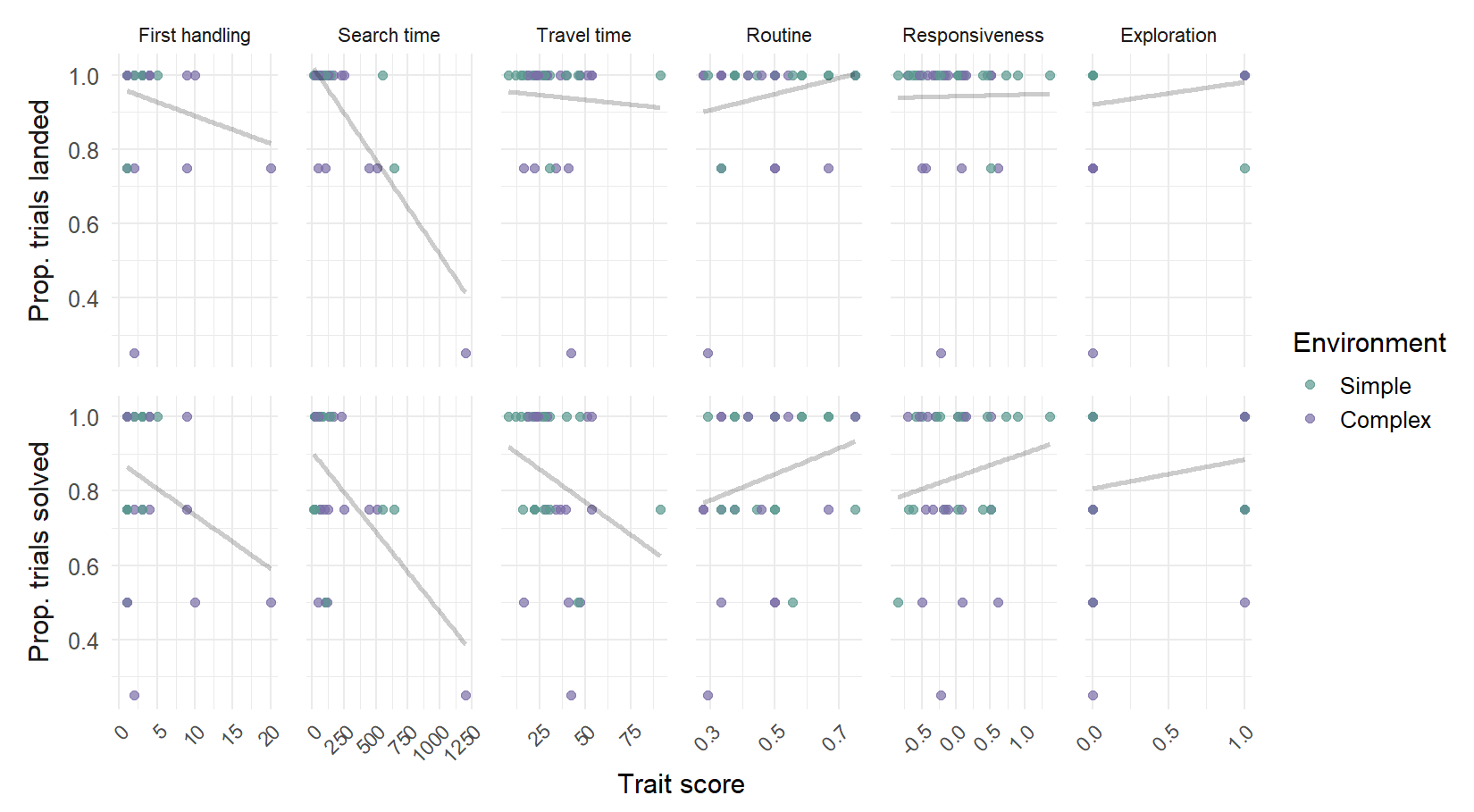

#### Supplementary Tables

###### **Table S1**: Effect of environment and trial on probability of landing on a novel flower.

|  | **Effect on landing on novel flower** | | |
| --- | --- | --- | --- |
| *Predictors* | *Odds Ratios* | *CI* | *p* |
| Intercept | 33.82 | 6.30 – 646.19 | **0.001** |
| Environment (complex vs simple) | 0.11 | 0.01 – 0.69 | **0.048** |
| Trial (Folded vs Bumpy) | 4.68 | 0.62 – 96.27 | 0.185 |
| Trial (Cap1 vs Bumpy) | 39475747.11 | 0.00 – NA | 0.992 |
| Trial (Cap2 vs Bumpy) | 1.41 | 0.27 – 8.01 | 0.681 |
| Observations | 140 | | |
| R^2^ Tjur | 0.093 | | |

###### **Table S2**: Effect of environment and trial on probability of solving a novel flower.

| **Effect on solving novel flower** | | |  |
| --- | --- | --- | --- |
| *Predictors* | *Odds Ratios* | *CI* | *p* |
| Intercept | 35.79 | 6.87 – 668.15 | **0.001** |
| Environment (complex vs simple) | 0.70 | 0.22 – 2.12 | 0.525 |
| Trial (Folded vs Bumpy) | 0.35 | 0.02 – 2.88 | 0.370 |
| Trial (Cap1 vs Bumpy) | 0.26 | 0.01 – 1.89 | 0.242 |
| Trial (Cap2 vs Bumpy) | 0.12 | 0.01 – 0.73 | 0.053 |
| Observations | 132 | | |
| R^2^ Tjur | 0.044 | | |

###### **Table S3**: Effect of environment and trial (and their interaction) on the time to solving a novel flower. Note that the response variable is log(time).

|  | **Effect on time to solve (reach reward) on novel flower** | | |
| --- | --- | --- | --- |
| *Predictors* | *Estimates* | *CI* | *p* |
| Intercept | 1.26 | 0.91 – 1.61 | **<0.001** |
| Environment (complex vs simple) | 0.17 | -0.36 – 0.71 | 0.515 |
| Trial (Folded vs Bumpy) | -0.03 | -0.53 – 0.47 | 0.906 |
| Trial (Cap1 vs Bumpy) | 1.17 | 0.68 – 1.66 | **<0.001** |
| Trial (Cap2 vs Bumpy) | 1.22 | 0.69 – 1.75 | **<0.001** |
| Env x Trial (Folded) | -0.50 | -1.24 – 0.24 | 0.182 |
| Env x Trial (Cap1) | -0.03 | -0.78 – 0.71 | 0.927 |
| Env x Trial (Cap2) | -0.19 | -0.97 – 0.60 | 0.638 |
| Observations | 117 | | |
| R^2^ / R^2^ adjusted | 0.466 / 0.432 | | |

###### **Table S4**: Effect of environment and trial on the time to abandoning a flower, i.e. the time from landing to leaving the flower for bees who did not manage to access reward. Note that because of the very small sample size, we did not include interactions. Here, the response variable is time (not log(time) ). We only include this result for completeness.

|  | **Effect on the time to abandon a flower without solving** | | |
| --- | --- | --- | --- |
| *Predictors* | *Estimates* | *CI* | *p* |
| Intercept | -0.14 | -39.84 – 39.56 | 0.994 |
| Environment (complex vs simple) | 2.68 | -19.88 – 25.25 | 0.796 |
| Trial (Folded vs Bumpy) | 39.44 | 2.92 – 75.96 | **0.037** |
| Trial (Cap1 vs Bumpy) | 38.44 | -0.02 – 76.90 | 0.050 |
| Trial (Cap2 vs Bumpy) | 15.93 | -24.68 – 56.53 | 0.403 |
| Observations | 15 | | |
| R^2^ / R^2^ adjusted | 0.519 / 0.327 | | |

###### **Table S5**: Effect of handling time in the first flower of the first trial (trial 1) on ‘innovation’, i.e. handling time (=solving time) on novel complex flowers (response variable is log(time) ).

|  | **Effect on solving time** | | |
| --- | --- | --- | --- |
| *Predictors* | *Estimates* | *CI* | *p* |
| Intercept | 1.15 | 0.83 – 1.47 | **<0.001** |
| First handling time | 0.31 | -0.01 – 0.63 | 0.059 |
| Trial (Folded vs Bumpy) | -0.06 | -0.51 – 0.39 | 0.795 |
| Trial (Cap1 vs Bumpy) | 1.21 | 0.76 – 1.67 | **<0.001** |
| Trial (Cap2 vs Bumpy) | 1.04 | 0.55 – 1.53 | **<0.001** |
| First hand x Trial (Folded) | -0.34 | -0.79 – 0.10 | 0.132 |
| First hand x Trial (Cap1) | -0.11 | -0.57 – 0.35 | 0.640 |
| First hand x Trial (Cap2) | 0.10 | -0.38 – 0.58 | 0.677 |
| Observations | 117 | | |
| R^2^ / R^2^ adjusted | 0.502 / 0.471 | | |

###### **Table S6**: Effect of search time in trials 6 & 7 (the Bumpy & Folded flowers) on time to solving flowers in all trials (6, 7, 8a, 8b) (response variable is log(time) ).

|  | **Effect on solving time** | | |
| --- | --- | --- | --- |
| *Predictors* | *Estimates* | *CI* | *p* |
| Intercept | 0.77 | -0.43 – 1.97 | 0.206 |
| Search time on B&F | 0.14 | -0.15 – 0.43 | 0.342 |
| Trial (Folded vs Bumpy) | 0.37 | -1.24 – 1.98 | 0.646 |
| Trial (Cap1 vs Bumpy) | 1.49 | -0.05 – 3.03 | 0.058 |
| Trial (Cap2 vs Bumpy) | 1.84 | 0.14 – 3.55 | **0.034** |
| Search x Trial (Folded) | -0.16 | -0.54 – 0.22 | 0.416 |
| Search x Trial (Cap1) | -0.09 | -0.45 – 0.27 | 0.635 |
| Search x Trial (Cap2) | -0.17 | -0.57 – 0.22 | 0.392 |
| Observations | 117 | | |
| R^2^ / R^2^ adjusted | 0.461 / 0.426 | | |

###### **Table S7**: Effect of travel time (average over trials 2, 3, and 4) on ‘solving time’ (i.e. handling time in the innovation trials, 6, 7, 8a and 8b) (response variable is log(time) ).

|  | **Effect on solving time** | | |
| --- | --- | --- | --- |
| *Predictors* | *Estimates* | *CI* | *p* |
| Intercept | -0.69 | -2.39 – 1.01 | 0.422 |
| Travel in trial 2-4 | 0.61 | 0.11 – 1.12 | **0.018** |
| Trial (Folded vs Bumpy) | 3.03 | 0.62 – 5.45 | **0.014** |
| Trial (Cap1 vs Bumpy) | 2.66 | 0.22 – 5.09 | **0.033** |
| Trial (Cap2 vs Bumpy) | 4.25 | 1.63 – 6.88 | **0.002** |
| Travel x Trial (Folded) | -1.00 | -1.72 – -0.27 | **0.007** |
| Travel x Trial (Cap1) | -0.45 | -1.18 – 0.27 | 0.219 |
| Travel x Trial (Cap2) | -0.94 | -1.73 – -0.16 | **0.019** |
| Observations | 117 | | |
| R^2^ / R^2^ adjusted | 0.498 / 0.466 | | |

###### **Table S8**: Effect of routine formation (SRI index, see Methods) on solving time (response variable is log(time) ).

|  | **Effect on solving time** | | |
| --- | --- | --- | --- |
| *Predictors* | *Estimates* | *CI* | *p* |
| Intercept | 0.18 | -0.79 – 1.15 | 0.720 |
| SRI (Routine formation) | 2.39 | 0.46 – 4.31 | **0.016** |
| Trial (Folded vs Bumpy) | 0.56 | -0.77 – 1.90 | 0.406 |
| Trial (Cap1 vs Bumpy) | 2.46 | 1.14 – 3.77 | **<0.001** |
| Trial (Cap2 vs Bumpy) | 1.84 | 0.45 – 3.22 | **0.010** |
| SRI x Trial (Folded) | -1.69 | -4.34 – 0.96 | 0.208 |
| SRI x Trial (Cap1) | -2.68 | -5.30 – -0.06 | **0.045** |
| SRI x Trial (Cap2) | -1.41 | -4.17 – 1.36 | 0.316 |
| Observations | 117 | | |
| R^2^ / R^2^ adjusted | 0.491 / 0.458 | | |

###### **Table S9**: Effect of responsiveness (see Methods) on solving time (response variable is log(time) ).

|  | **Effect on solving time** | | |
| --- | --- | --- | --- |
| *Predictors* | *Estimates* | *CI* | *p* |
| Intercept | 1.32 | 1.06 – 1.58 | **<0.001** |
| Responsiveness | -0.39 | -0.88 – 0.11 | 0.125 |
| Trial (Folded vs Bumpy) | -0.25 | -0.62 – 0.11 | 0.176 |
| Trial (Cap1 vs Bumpy) | 1.16 | 0.80 – 1.53 | **<0.001** |
| Trial (Cap2 vs Bumpy) | 1.15 | 0.76 – 1.54 | **<0.001** |
| resp x Trial (Folded) | 0.72 | 0.01 – 1.43 | **0.047** |
| resp x Trial (Cap1) | 0.25 | -0.45 – 0.96 | 0.479 |
| resp x Trial (Cap2) | 0.64 | -0.12 – 1.41 | 0.099 |
| Observations | 117 | | |
| R^2^ / R^2^ adjusted | 0.479 / 0.446 | | |

###### **Table S10**: Effect of exploration (see Methods) on solving time (response variable is log(time) ).

|  | **Effect on solving time** | | |
| --- | --- | --- | --- |
| *Predictors* | *Estimates* | *CI* | *p* |
| Intercept | 1.23 | 0.91 – 1.56 | **<0.001** |
| Exploration | 0.28 | -0.26 – 0.82 | 0.309 |
| Trial (Folded vs Bumpy) | -0.00 | -0.47 – 0.46 | 0.996 |
| Trial (Cap1 vs Bumpy) | 1.21 | 0.74 – 1.67 | **<0.001** |
| Trial (Cap2 vs Bumpy) | 1.27 | 0.76 – 1.77 | **<0.001** |
| Expl x Trial (Folded) | -0.70 | -1.45 – 0.06 | 0.071 |
| Expl x Trial (Cap1) | -0.15 | -0.91 – 0.60 | 0.690 |
| Expl x Trial (Cap2) | -0.33 | -1.12 – 0.46 | 0.411 |
| Observations | 117 | | |
| R^2^ / R^2^ adjusted | 0.473 / 0.439 | | |
